# The Popgen Pipeline Platform: A Software Platform for Facilitating Population Genomic Analyses

**DOI:** 10.1101/785774

**Authors:** Andrew Webb, Jared Knoblauch, Nitesh Sabankar, Apeksha Sukesh Kallur, Jody Hey, Arun Sethuraman

## Abstract

Here we present the Pop-Gen Pipeline Platform (PPP), a software platform with the goal of reducing the computational expertise required for conducting population genomic analyses. The PPP was designed as a collection of scripts that facilitate common population genomic workflows in a consistent and standardized Python environment. Functions were developed to encompass entire workflows, including: input preparation, file format conversion, various population genomic analyses, output generation, and visualization. By facilitating entire workflows, the PPP offers several benefits to prospective end users - it reduces the need of redundant in-house software and scripts that would require development time and may be error-prone, or incorrect. The platform has also been developed with reproducibility and extensibility of analyses in mind. The PPP is an open-source package that is available for download and use at https://ppp.readthedocs.io/en/latest/PPP_pages/install.html

## Introduction

Since the advent of genomics, population genetics has quickly become dominated by complex statistical and computational methodologies [1, 2]. An unfortunate consequence of this fact is that many investigators lack the necessary resources - computational, and time - to independently implement many of these methodologies. This inevitably requires investigators to select from a plethora of software (i.e. analytical tools) that have been developed by other researchers. While this is not inherently a problem, and a common practice among many professions, it is not without its own difficulties. Investigators frequently face bespoke input and output formats that may not be accompanied by an intuitive and easy-to-use file-format conversion software, implementations that may be complex and open to misinterpretation, and lastly implementations incapable of large-scale analyses.These challenges are further amplified as few analyses require a single tool, but rather require an analytical pipeline. Analytical pipelines typically incorporate a number of methodologies and software designed specifically to connect those methodologies in a specific order.

The challenges posed by analytical pipelines have been partially mitigated by the development of software packages or “tool-kits” that provide tools for a variety of methodologies. However, while popular packages such as `vcftools` [3], `bcftools` [4], and `plink` [5] have proven invaluable to many investigators, they cannot be all-encompassing. The absence of such tool-kits often requires investigators, if able, to create pipelines that are frequently recreated, infrequently published, time consuming to develop, and susceptible to error. For these reasons, analyses based on such pipelines are often difficult or impossible to completely replicate [6, 7], which is an issue of growing concern in research [8].

In an attempt to greatly alleviate these obstacles we have developed the PopGen Pipeline Platform (PPP). The PPP was designed to be a comprehensive platform wherein investigators can conduct many of the analytical pipelines involved in population genomics in a simple and standardized environment. We achieved this goal by incorporating and connecting various tool-kits, standard tools/methods, and common analytical practices. To demonstrate both the simplicity and the comprehensive nature of the PPP, we designed and implemented population genomic analyses of publicly available data from chimpanzees [9] using only the PPP.

## New Approach

### Design

The PPP was written in the Python programming language and designed to operate using either Python versions 2 or 3. Python was selected primarily to reduce the complexity of future development, take advantage of various relevant and powerful Python libraries, and to minimize compatibility issues for prospective users. The PPP was designed as a collection of modular functions that may be combined to offer a wide variety of analyses and pipelines required by population geneticists. The core functions of the PPP - i.e. functions commonly used among analyses - were designed to operate using VCF-based file formats [3]. This decision was due to the predominance of the VCF file format within the population genomics community, specifically the frequent support for this format among tools, and the likelihood of most publicly available datasets being made available as VCF formatted files. Most hypothetical runs in the PPP will begin with these core functions, and then branch off into the desired combination of analysis-specific functions. It should be stated that most analysis-specific functions do not support VCF-based file formats, but rather incorporate a preceding file conversion core function to operate. This design was chosen to avoid superfluous conversions, many of which are computationally intensive.

A fundamental aspect of the PPP’s design is that if a specific technique (e.g. tool, software package, statistic) is synonymous with an analysis, that technique will be integrated into the function associated with the analysis. In some instances we have integrated multiple techniques into a single function - e.g. we have included both `BEAGLE` [10] and `SHAPEIT` [11] in our phasing function. As prospective users may not be familiar with a technique, relevant information and links to the original material may be found within the documentation and appropriate references will be provided upon use of a technique.

The PPP was also designed to include other features to further simplify and expedite analyses. For instance, the PPP integrates a versatile configuration system that allows prospective users to configure functions in two ways: with optional command-line arguments; or with optional arguments specified within a configuration file. By using a configuration file it is possible for prospective users to configure an entire analysis or pipeline. This is possible due to the standardized argument scheme designed for the PPP which allows the assignment of global arguments - i.e. consistent among the entire platform - and function-specific arguments - such as the explicit input and output for each function. Another feature of the PPP is the use of the Model file format that we developed for use in the platform. The Model file is a JSON-based format that is able to store multiple population models, including the relevant details of each model (i.e. populations, individuals, population tree, and other relevant meta-data). A primary benefit of the Model file is the ability to automatically assign information from the specified model to functions, such as the populations and their associated individuals. The file also simplifies record keeping as it becomes the repository for model-related information.

### Overview

A consequence of the design of the PPP is that a hypothetical analysis could use a combination of functions that do not demonstrate the comprehensive nature of the platform - see Figure 1. for an illustration of the initial release of the PPP. Therefore, to give a sufficient overview of the PPP, we have chosen to describe the functions required in the Isolation with Migration (IM) [12] pipeline we used for analyzing population genomic data from chimpanzees [9]. As the demographic history of the chimpanzees have been extensively studied [13, 14, 15, 16], we selected two closely related populations - Central chimpanzees (*Pan troglodytes troglodytes*) and Western chimpanzees (*Pan troglodytes verus*) - to demonstrate the effectiveness of the PPP in comparison to similar analyses. In particular, we wished to explore the divergence of the two populations by estimating their population sizes, migration rates, and divergence time using multi-locus genomic data under an IM model.

**Figure 1:**
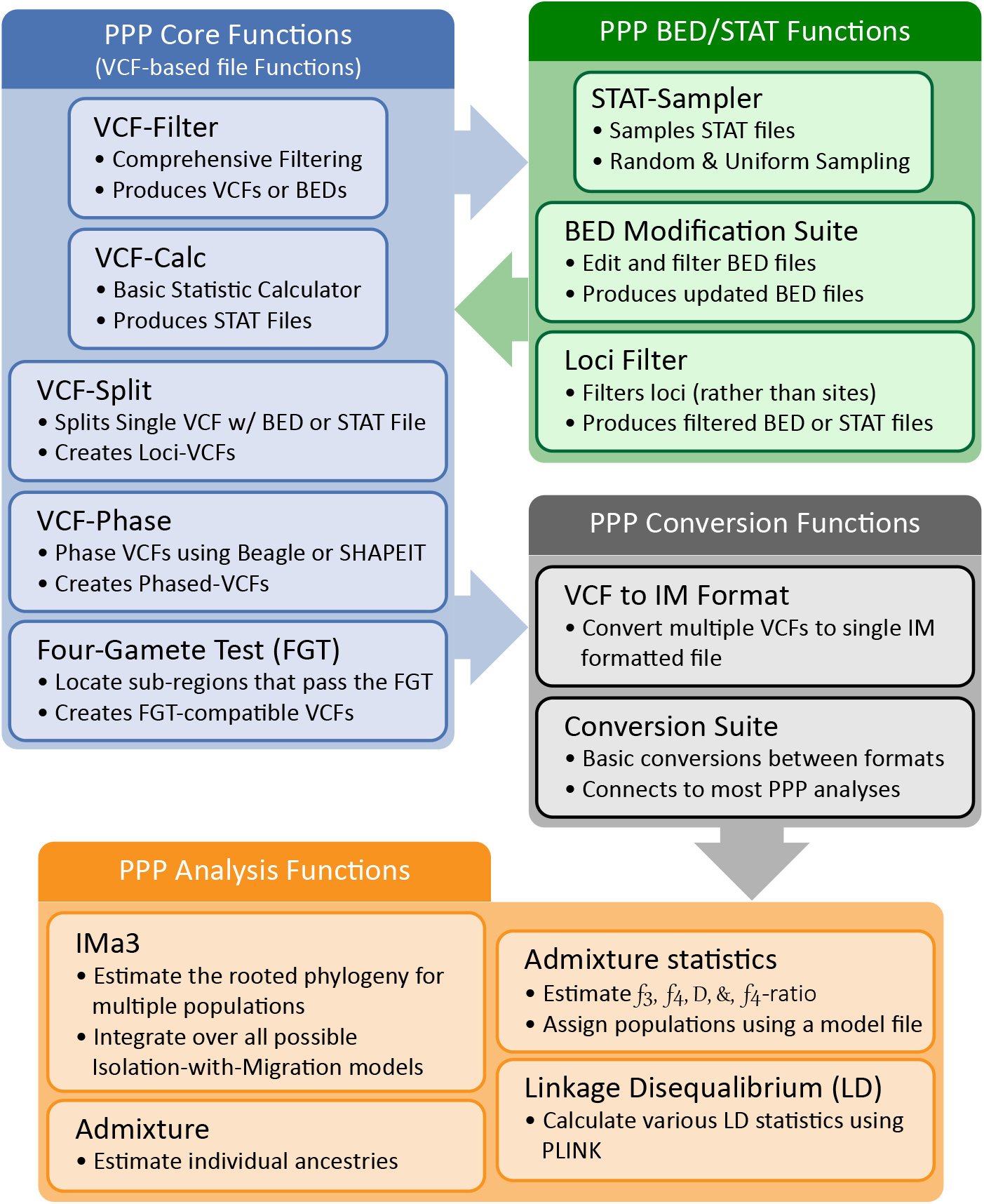
Structure of the PPP. PPP functions are grouped into four categories: i) the Core PPP functions that operate on VCF files; ii) the optional BED and STAT functions which may be used to sample, filter, and/or edit BED or STAT files; iii) the conversion functions which are required to convert from VCF to analysis-specific file formats; and iv) the analysis functions which are used to automate their respective analyses.

The first procedure in our analysis pipeline was applying filters to remove sites with missing data and non-biallelic sites. The removal of non-biallelic sites (i.e. multiallelic sites) is of particular importance as they violate the Infinite Sites (IS) model [17] assumption of a single polymorphism per site assumed by the IM model [12] implemented in our analysis pipeline. It also bears mentioning that additional downstream procedures are also required to avoid other violations of model assumptions, and will be reported where relevant. The filter procedure of our analysis was completed using the PPP’s **VCF-filter** function. **VCF-filter** was designed to perform filtering operations on VCF-based files and is expected to be the first function in most analyses. Prospective users are able to select from a comprehensive collection of filters that are assigned alongside a specified value (i.e. a threshold) or a specified file (e.g. a BED file containing genomic coordinates of sites to remove, or include) (Table 1). Depending on the needs of the prospective user, the function is capable of returning either a BED file of the filtered sites, or an updated VCF-based file. If a model is specified from a Model file, the function is designed to automatically remove non-relevant samples before filters are applied.

**Table 1:** Capabilities of the PPP Filters and Statistic Calculator.

| Function | Purpose | Capabilities |
| --- | --- | --- |
| VCF-filter | Filtering Variants | *Include/exclude variants sites by:* genomic position, missing data count and percentage, allele count, MAF, MAC, presence of indels, SNP IDs, associated with a specific flag (i.e. PASS) |
| VCF-calc | Statistic Calculator | *Calculate the following statistics:* $F_{st}$ (site- and window-based), Tajima's $D$, Nucleotide Diversity (site- and window-based), allele frequency, inbreeding coefficients ($F_{IT}$ and $F_{IS}$), tests of Hardy-Weinberg Equilibrium |
| informative-loci-filter | Filtering Loci | *Include/exclude loci by:* informative site count, variant site count, missing data count, ignoring indels, ignoring multiallelic variants, ignoring CpG sites |

Our analysis pipeline then proceeded to randomly sample 200, 1 kbps, non-overlapping genomic loci by their respective *F*_*st*_ values between the two populations. Due to subsequent analytical requirements, and the assumption of putatively neutral sites [12], each sampled locus was required to be both informative - i.e. in possession of at least four variant positions - and intergenic. This procedure, **VCF-calc** to calculate the *F*_*st*_ values within 1 kbps loci, **informative-filter** to remove loci that were not informative, and **stat-sampler** to pseudo-randomly sample 200 loci. **VCF-calc** was designed to calculate many of the basic statistics used in population genetic analyses on VCF-based files (Table 1). For most statistics, little to no configuration is necessary, however, some statistics do require additional parameters (e.g. window length, window step length) to operate. This function is designed to return a tabular statistic output file that is usable by other functions within the PPP. If a model is specified from a Model file, the function is designed to automatically assign the relevant populations and/or individuals to compute the appropriate statistics. The **informative-filter** function was designed to apply various locus-based filters often required by population genomic analyses on VCF-based files (Table 1). In comparison to **VCF-filter**, these filters evaluate and filter each locus as a single entity. To operate, the majority of filters only require a BED or statistic file to define the loci of interest. Filters were also designed to be easily configurable by altering default values or by enabling optional parameters. **informative-filter** is designed to return a filtered copy of the original BED or statistic file. **stat-sampler** was designed to pseudo-randomly sample loci from statistic files produced by the **VCF-calc** function. Prospective users may select from one of two pseudo-random sampling schemes: a random scheme that samples loci from the entire file, and a uniform sampling scheme that samples loci from equally sized bins derived from the statistic of choice. **stat-sampler** may also be configured to alter both sampling schemes - e.g. samples to select and number of bins - and to reproduce previous results, if desired. The function is designed to return a sampled version of the statistic file as output.

The next procedure in our analysis pipeline was the creation of phased VCF-based files for each of the sampled loci. Phased chromosomes are required for our pipeline to identify potential recombination events by the Four-gamete Test [18]. It should be noted that phasing was possible prior to the creation of individual VCF-based files for each sampled locus, but is computationally demanding. Our procedure required the use of the **VCF-split** function to generate locus-specific VCF-based files and **VCF-phaser** function to phase the files. The **VCF-split** function was designed to split a single VCF-based file using either a BED or statistic file to define the coordinates for the loci of interest. If a model is specified from a Model file, the function is designed to only return the relevant individuals in the loci VCF-based files. **VCF-phaser** was designed to phase VCF-based files using either `SHAPEIT` [11] or `BEAGLE` [10]. Phasing with **VCF-phaser** only requires prospective users to specify a VCF-based file - which by default uses `SHAPEIT` [10]. However, **VCF-phaser** may be configured to instead phase VCF-based files with `BEAGLE` [10] or configure the settings of either algorithm. If a model is specified from a Model file, the function is designed to only phase and return the relevant individuals.

Our pipeline next required the identification of sub-regions of each locus without recombination within our phased VCF-based files. This procedure was necessary to avoid violating the assumption of no recombination within loci of the IM model [19]. This was accomplished using the **Four-gamete Test** function of the PPP, which was designed to check for the presence of recombination events between pairs of segregating sites [18]. The PPP’s implementation of the **Four-gamete Test** takes a VCF-based file of a kilobase-scale region in a chromosome, then finds sub-regions of the loci that have less than four gametes among them. Prospective users may configure the **Four-gamete Test** to: require a specific number of informative sites; return either a single or all compatible sub-regions; ignore multiallelic sites; and include sites with missing data. By default, the function is designed output a VCF file of a sub-region with at least two informative sites that passed the test.

The last procedure in our pipeline was performing an IM analysis using `IMa3` [16]. However, before we were able to proceed to the IM analysis of our pipeline we were required to convert the sub-region VCF-based files into a single IM formatted file that is compatible with our implementation of IMa3 [16]. This procedure was accomplished using the **vcf-to-ima** conversion function of the PPP. **vcf-to-ima** was designed to automatically generate an IM formatted file from a collection of sub-region VCF-based files, a model specified from a Model file, and additional parameters provided by the prospective user. This design allows for IM formatted files to be easily configured by specifying a different Model or altering parameters. Once the conversion process was finished we used the PPP function **ima3-wrapper** to perform all IM analyses. **ima3wrapper** handles the passing of input parameters to IMa3, while also handling multi-threading in the subprocess calls if the user specifies. Most required input is specified in the IM input file, with additional options required to specify upper limits, priors for parameters to be estimated, and determine how long to burn-in, and genealogy sampling run-time of the MCMC should be. The final output is a file with estimates of population model parameters (migration rates, population sizes, and divergence times), with confidence intervals around these estimates.

Finally, while our pipeline focused on performing an IM analysis, the PPP was designed to easily allow the implementation of additional analyses, if desired. For example, we could use many of the files produced in our IM analysis to estimate population structure using ADMIXTURE [20], test for introgression using AdmixTools [21], or linkage disequilibrium using PLINK [5].

## Results

To demonstrate the capabilities of the PPP we compared an Isolation with Migration analysis of two chimpanzees populations to previous reports [13, 14]. We found our estimates of the divergence time, the ancestral chimpanzee population size, migration rates, and the populations sizes of the extant chimpanzee populations - central chimpanzees (*Pan troglodytes troglodytes*) and western chimpanzees (*Pan troglodytes verus*) to be consistent with previous findings (Table 2).

**Table 2:** Evolutionary history of Central and Western Chimpanzees, estimated using PPP and IMa3. The mean and highest posterior parameter estimated population sizes (q), migration rates (m), and divergence time (t) between *P. t. troglodytes* (population 0) and *P. t. verus* (population 1).

| Parameter | Mean | Highest Posterior |
| --- | --- | --- |
| q0 | 1.219 | 1.204 |
| q1 | 0.3469 | 0.3400 |
| q2 | 0.7531 | 0.7640 |
| $m0 \rightarrow m1$ | 0.5860 | 0.5675 |
| $m1 \rightarrow m0$ | 0.8330 | 0.7925 |
| t | 0.4155 | 0.4494 |

## Discussion

The primary goal behind the development of the PPP was to create a simple, standardized, and robust platform for population genetic analyses. Ideally, an end user would only require a specific combination of PPP functions to implement their desired pipeline. To demonstrate this capability, we examined the demographic history of two closely related chimpanzee population and compared the results to previous findings [14, 15, 16]. We found that the PPP greatly reduced the overall complexity of our analysis and was able to successfully reproduce previous findings. With the exception of downloading the necessary files (e.g. chimpanzee VCF input, BED files containing gene coordinates) all operations were completed using PPP functions alone. Assembling the pipeline was a straightforward process as the majority of functions could be invoked in tandem without requiring intermediate processing steps. We were also able to quickly process the VCF input for our IM analysis as the majority of PPP functions required less than 5 minutes to operate, with the exception being the initial filtering procedure which took roughly 50 minutes and the IM analysis which required approximately 400 hours of CPU time. We also found that repeating our analysis - either to explore the results of different parameters, reproduce our findings, or remedy errors - was a simple process and could be done rapidly if the initial filtering was not repeated. Taken together, the PPP has achieved its primary design goal, but that does not signify the platform is complete. Additionally, this sample pipeline, along with other examples have been published as Jupyter Notebooks on the PPP’s development website.

Future development of the PPP will primarily be focused on improvements to the platform. First and foremost is the creation of a Galaxy Project [22] wrapper to expand the user base of the platform, primarily to assist users more familiar with a graphical user interface and/or web applications. As the PPP was developed in consideration of an eventual Galaxy wrapper, implementing this improvement will be straightforward. We also intend to have ongoing releases of additional population genetic analyses for the platform. The modular structure of the platform should allow for the majority of these updates to only require creation of the function to automate the analysis and potentially updating the file conversion suite. Future releases will also focus on improvements to the overall speed (and efficiency) of the platform. One potential improvement currently being explored is the incorporation of Cython, which aims to achieve C-like performance among python scripts [23]. We also plan on exploring the possibility of using Jupyter notebooks [24] to store and share analysis pipelines. Jupyter notebooks are a simple and ideal format for analysis pipelines as they allow computer code - e.g. a PPP function - to be accompanied by textual elements, such as descriptions of each function and their overall purpose.

## Acknowledgments

This work was supported by an NSF ABI Grant 1564659 to AS and JH. This research includes calculations carried out on Temple University’s HPC resources and thus was supported in part by the National Science Foundation through major research instrumentation grant number 1625061 and by the US Army Research Laboratory under contract number W911NF-16-2-0189.

## Notes

https://ppp.readthedocs.io/en/latest/PPP_pages/install.html

https://www.csusm.edu/ppp/

